# SEN virus genotype H distribution in β-thalassemic patients and in healthy donors in Iraq: Molecular and physiological study

**DOI:** 10.1101/821744

**Authors:** Mushtak T.S. Al-Ouqaili, Yasin H. Majeed, Sahar K. Al-Ani

## Abstract

The SEN virus (SENV) has been linked to transfusion-associated non-A-E hepatitis; however, information regarding SENV infections in patients with thalassemia, particularly in those with hepatitis virus co-infections, remains limited. This study investigated the frequency of SENV (genotypes D and H) infections in Iraqi patients with thalassemia who were and were not infected with hepatitis C virus (HCV). The study involved 150 β-thalessemia patients (75 with HCV infections and 75 without) and 75 healthy blood donors. Patient levels of vitamins C and E, liver function markers, and glutathione peroxidase (GPX) were determined. Recovered viral nucleic acids were amplified using the conventional polymerase chain reaction (SENV DNA) or the real-time polymerase chain reaction (HCV RNA) techniques. Only 10% of healthy donors had evidence of SENV infection. Among patients with thalassemia, 80% and 77% of patients with and without concurrent HCV infections, respectively, had SENV infections. DNA sequencing analyses were performed on blood samples obtained from 29 patients. Patients with thalassemia, particularly those with SENV infections, had higher levels of several enzymatic liver function markers and total serum bilirubin (P < 0.05) than did healthy blood donors. Among the examined liver function markers, only gamma-glutamyl transferase demonstrated significantly higher levels in HCV-negative patients infected with SENV-H than in those infected with SENV-D (P = 0.01). There were significantly lower vitamin C, vitamin E, and glutathione peroxidase levels in patients than in healthy donors (P < 0.05), but only glutathione peroxidase levels were significantly lower in HCV-negative thalassemia patients infected with SENV than in those without SENV infections (P = 0.04). The SENV-H genotype sequences were similar to the global standard genes in GenBank. These results increase our understanding of the nature of the SENV-H genotype and the differential role of SENV-H infections, compared to SENV-D infections, in patients with thalassemia, in Iraq.

**Author summary:** In patients with β-thalassemia, regular blood transfusions increase patient survival but increase the risk of acquiring blood-borne viral infections, especially viral hepatitis. The SEN virus (SENV) is associated with non-A-E hepatitis but its exact role in the pathogenesis of liver disease remains unknown. This study investigated the frequency of SENV infections among Iraqi patients with thalassemia, with and without hepatitis C infections. The study revealed that the prevalence of SENV infections is significantly higher in patients with β-thalassemia, regardless of their hepatitis C infection status, than in a healthy population of blood donors. The two most common genotypes of the virus (D and H) have generally similar physiological impacts as both increase the levels of markers of hepatic dysfunction in thalassemia patients. However, SENV-H infections were associated with significantly higher levels of gamma-glutamyl transferase in HCV-negative patients with thalassemia, potentially predicting hepatocellular carcinoma development. Although thalassemia patients demonstrated significantly lower levels of the antioxidants vitamins C and E, compared with healthy donors, only the levels of glutathione peroxidase (another antioxidant) were significantly lower in SENV-infected patients than in non-SENV-infected patients. This study aids our understanding of the differential effects of SENV-D and SENV-H infections in β-thalassemia patients.

## Introduction

Thalassemia, a common hereditary hemoglobinopathy, is characterized by the abnormally low production of hemoglobin. The low levels of hemoglobin result in anemia, which necessitates treatment with frequent blood transfusions. In Iraq, the prevalence of thalassemia increased slightly between 2010 and 2015, from 33.5/100,000 to 37.1/100,000, whereas the incidence of the disease decreased, from 72.4/100,000 live births to 34.6/100,000 live births, during the same period [1]. As a result of the regular transfusions required by individuals with thalassemia, many of these individuals acquire blood-borne infections. In Iraq, the same study that investigated the incidence and prevalence of thalassemia determined that patients were often infected with hepatitis C virus (HCV, 13.5%) or hepatitis B virus (0.4%) at some point during their lives [1].

Viral hepatitis is a significant cause of morbidity and mortality, worldwide, particularly in the tropics. It is caused by at least five distinct viruses, each with unique molecular characteristics and replication cycles but sharing a common tropism for the liver and causing overlapping clinical patterns of disease [2]. The hepatitis viruses are the most common chronic blood-borne viruses associated with this disease, but other infectious agents have been suggested to cause viral hepatitis that are not directly attributed to the hepatitis viruses (non-A–E hepatitis). The latest virus suggested to have a role in non-A–E hepatitis is the SEN virus (SENV). This virus, discovered in 1999 in the blood of a human immunodeficiency virus-infected patient, is a 26-nm, single-stranded DNA virus that is distantly related to the TT virus [3]. Phylogenetic analyses identified eight different SENV strains belonging to the *Circoviridae* family, a group of small DNA viruses that includes the TT virus [4]. Of the 9 SENV genotypes identified, to date, two (SENV-D and SENV-H) have been extensively studied and are present in approximately 2% of blood donors in the USA, 2% of donors in Italy, and 10% of donors in Japan; they appear to be readily transmitted by blood transfusions and other common parenteral routes [1].

SENV infections, particularly those caused by the D and H genotypes, are frequently associated with non-A–E hepatitis, giving rise to the suggestion that the virus may be the causative agent. However, there is no firm evidence of the virus causing hepatitis or worsening existing disease [5,6]. Co-infections involving SENV and the hepatitis C virus (HCV) or hepatitis B virus (HBV) are also common because the viruses share similar transmission routes (e.g., blood transfusions) [7]. Our previous study [8], demonstrated the presence of SENV viremia in Iraqi patients with β-thalassemia who were and were not infected with HCV. Further, that study also demonstrated that both the D and H SENV genotypes were present in Iraqi patients and noted the impact of the viral infection on liver function indicators in infected individuals. As the previous study was relatively small, this study sought to confirm the previous results and to further investigate the potential for differential effects caused by the SENV D and H genotypes on patients with β-thalassemia.

## Patients and methods

A total of 150 patients with β-thalassemia were referred to the Hereditary Blood Disease Center (Baghdad, Iraq) and included in this retrospective study. These individuals were divided into two groups, according to their HCV infection status; equal numbers (n = 75) were HCV-positive and HCV-negative. Another 75 individuals were randomly recruited into the study from among the healthy blood donors attending the Iraqi National Center of Blood Transfusion. The healthy donors were afebrile, not jaundiced, did not demonstrate any signs of chronic liver disease; further, none had any known contact with individuals with jaundice. The ages of included participants ranged from 5 to 44 (mean, 18 ± 7.8) years and all received regular blood transfusions or donated blood between January and May 2018. All procedures performed in this study involving human participants were in accordance with the University of Anbar Ethical Approval Committee (authentication no. 31, December 6, 2017). Because of the retrospective nature of this study and the use of anonymized data from samples collected for clinical purposes, the need for informed consent was waived.

## Serology

### Liver function markers

A blood chemistry analyzer (Celercare^®^ M1, MNCHIP, Tianjin, China) and lyophilized liver function panel kits (MNCHIP) were used to determine levels of alanine transaminase (ALT), aspartate aminotransaminase (ASP), alkaline phosphatase (ALP), total serum bilirubin (TSB), and gamma-glutamyl transferase (GGT).

### Detection of anti-HCV-antibodies

All serum samples were examined for the presence of anti-HCV antibodies using enzyme-linked immunosorbent assay (ELISA) kits (Human Gesellschaft fur Biochemica und Diagnostica, Wiesbaden, Germany).

### Vitamins C and E and glutathione peroxidase (GPX) levels

ELISA-based double antibody sandwich kits (Shanghai Yehua Biological Technology, Shanghai, China) were used to assay levels of human vitamin C (catalog no. YHB3202Hu), vitamin E (catalog no. YHB3208Hu), and GPX (catalog no. YHB1369Hu) in patient serum samples.

## Molecular study

### Nucleic acid extraction

Viral nucleic acid extraction kits (SaMag Viral Nucleic Acids Extraction Kit, Sacace Biotechnologies, Como, Italy) were used to extract SENV DNA and HCV RNA from participant plasma specimens. Briefly, the extraction process involved lysing, binding, washing, and eluting the nucleic acids. Frozen plasma samples were thawed at room temperature (15–25°C) and processed in an automated instrument (SaMag-12, Sacace Biotechnologies), after equilibrating to room temperature [8].

### Nucleic acid concentration and purity

A quantum fluorometer (Promega, Madison, WI, USA) was utilized to measure the concentration of extracted nucleic acids; preprogrammed settings were available for quantitating DNA (QuantiFluor^®^ ds DNA, Promega) and RNA (QuantiFluor^®^ RNA, Promega). A volume (100 μL) of diluted nucleic acid sample (1 μL of nucleic acid + 99 μL buffer) was mixed with the appropriate QuantiFluor dye. After 5 minutes of incubation at room temperature, the DNA or RNA concentrations were determined in the fluorometer. The nucleic acid samples were also read in a spectrophotometer, equipped with Nanodrop software (ThermoFisher, Waltham, MA, USA), at 260 nm and 280 nm. If the results were between 1.7 and 1.9, the samples were considered to be contamination free and adequate for further analyses.

### HCV real-time polymerase chain reaction (PCR) quantification

HCV Real™ Quant kits (Sacace Biotechnologies) were used for the quantitative detection of HCV in human plasma. Briefly, HCV RNA was extracted from plasma samples, amplified, and detected using fluorescent reporter dye probes specific for either HCV or immune complexed HCV. An internal control served as the extraction and amplification control for each individually processed specimen, allowing identification of possible inhibitions. Immune complexes were detected in a specifically labeled channel other than that used for HCV RNA. Real-time monitoring of fluorescence intensities allowed detection and quantification of the accumulating product without opening the reaction tube after real-time amplification. The internal control was detected on the FAM channel and HCV RNA on the CY3 channel. For each control and patient specimen, the concentration of HCV RNA was calculated using the following formula: HCV RNA copies/specimen (CY3 channel)/immune complexed RNA copies/specimen (FAM channel) × coefficient = IU HCV/mL. These results were also expressed as copies/mL by multiplying by 4 [9, 10].

### PCR amplification of SENV DNA

The primers used in the study were SENV-AI-IF (5’TACTCCAACGACCAGCTAGACCT3’), SENV-AI-IR (5’GTTTGTGGTGAGCAGAACGGA3’), SENV-D-1148 F (5’GCAGTTGACCGCAAAGTTACAAGAG3’), and SENV-D-1341 R (5’GCAGTTGACCGCAAAGTTACAAGAG3’) (AlphaDNA, Montreal, QC, Canada). The template DNA and primers were added to a PCR tube, along with nuclease-free water, to a total volume of 50 μL. The PCR reaction was carried out in a 50-μL mixture containing *Taq* DNA polymerase, dATP, dGTP, dCTP, dTTP, 1.5 mM MgCl_2_, reaction buffer (pH 9), loading dye buffer (yellow and blue dyes), amplification primers (2 μL, each), target DNA (10 μL), and nuclease-free water [8].

The SENV DNA was amplified using nested conventional PCR, as follows. The first-round amplification was: 94° C for 4 min, 1 cycle for initial denaturation of the template; 35 cycles at 94° C for 40 s to denature the DNA; 55° C for 50 s, 35 cycles for annealing; 72° C for 50 s, 35 cycles for extension; finally, 72° C for 10 min, 1 cycle for final extension. The second PCR run was conducted as follows: 94° C for 4 min, 1 cycle for initial denaturation of the template; 35 cycles at 94° C for 30 s to denature the nucleic acid; 55° C for 50 s, 35 cycles for annealing; 72° C for 50 s, 35 cycles for extension; and 72° C for 10 min, 1 cycle for final extension.

Upon PCR completion, 10 μL of amplified DNA was mixed with 4 μL of Redsafe nucleic acid stain and loaded on to 2% agarose gels. After electrophoresis was complete, the gel was placed on an ultraviolet transilluminator and digitally documented.

### Sequencing

The National Instrumentation Center for Environmental Management (Seoul, Korea) sequenced the PCR products on our behalf. The sequences were then run through the standard gene BLAST program (http://www.ncbi.nlm.nih.gov) and sequences were aligned using the BioEdit program (Ibis Therapeutics, Carlsbad, CA, USA). An evolutionary analysis was conducted using Molecular Evolutionary Genetics Analysis software (version 6, Pennsylvania State University, State College, PA, USA) [11].

### Statistical analysis

Data are presented as percentages, means, standard deviations, and ranges. Significant differences between means (quantitative data) were examined using Students *t*-test for differences between two independent means, paired *t*-tests for differences between paired observations (or two dependent means), or analyses of variance for differences among more than two independent means. Significant differences between percentages (qualitative data) were determined using the Chi-square test, applying Yate’s correction or Fisher’s Exact test when applicable. All statistical analyses were performed using SPSS (ver. 22, IBM, Armonk, NY, USA); a P-value ≤ 0.05 was considered significant.

## Results

The 75 patients with β-thalassemia and strongly suspected of having HCV infections were confirmed to be HCV-positive; the other 75 patients with thalassemia and the 75 healthy donors were confirmed to be HCV-negative. Nested conventional PCR was used to identify SENV infections, with the results revealing a significantly greater occurrence of SENV infections among patients with thalassemia than among healthy donors. Agarose gel electrophoresis effectively differentiated between the 193-bp SENV-D and 118-bp SENV-H genotypes as well as demonstrated their co-occurrence in some individuals (Figs 1 and 2).

**Fig 1.**
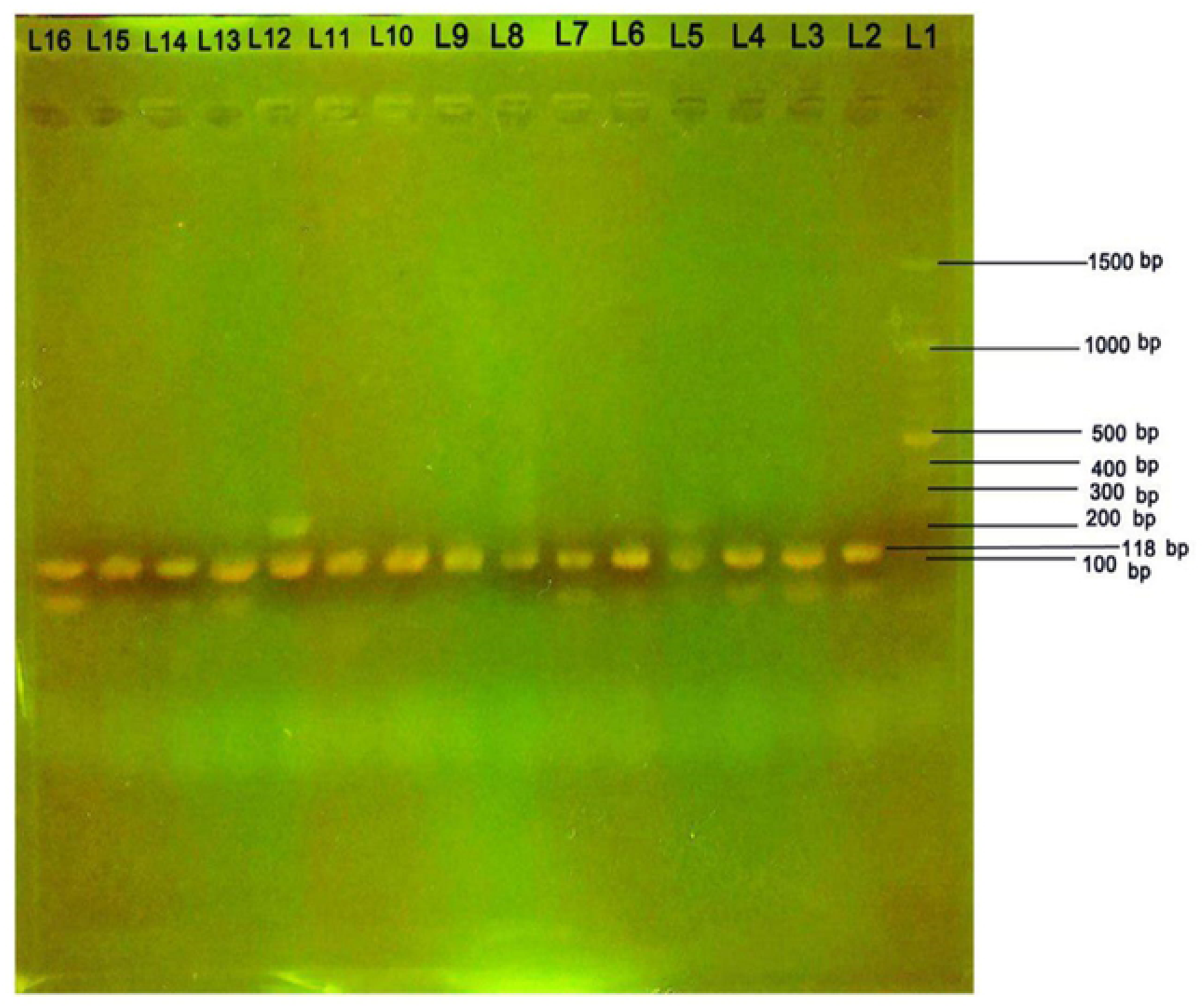
Agarose gel electrophoresis of amplified SEN virus H-gene DNA. Bands showing the amplified SEN virus-H gene (118 bp) are shown in lanes L2–L15. DNA molecular weight markers (100–1500 bp) are present in lane L1.

**Fig 2.**
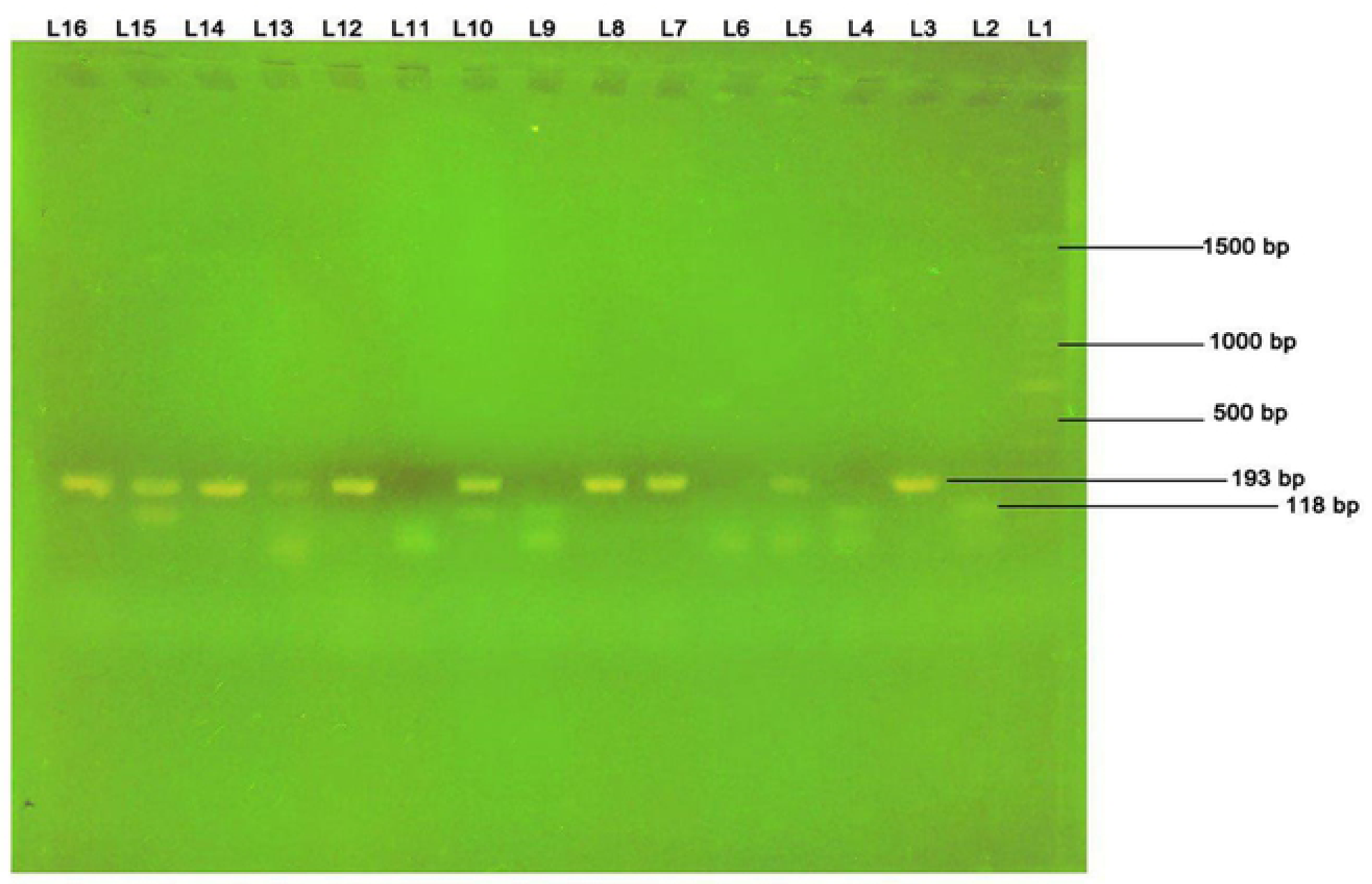
Agarose gel electrophoresis of amplified SEN virus D- and H-gene DNA. Bands showing the presence of SEN virus D (193 bp) and H (118 bp) genes in lanes L5, L10, and L15. DNA molecular weight markers (100–1500 bp) are present in lane L1.

Table 1 shows that the frequency of SENV infections among patients with thalassemia (78%) was significantly higher than among healthy blood donors (10%; P < 0.05). Table 1 also emphasizes that the prevalence of SENV infections in patients with thalassemia was not dependent on their HCV infection status. Interestingly, among the healthy blood donors with SENV infections, more SENV-positive individuals lived in urban environments (9/51, 17.6%) than lived in rural environments (1/24, 4.1%); this was a statistically significant difference (p = 0.003).

**Table 1.** SEN virus infection rates among healthy blood donors and among patients with β-thalassemia, with and without evidence of hepatitis C virus (HCV) infection.

| Patients with β-thalassemia | Healthy<br>donors |
| --- | --- |

| SEN virus status | HCV RNA-positive<br>(n = 75) | HCV RNA-negative<br>(n = 75) | (n = 75) |
| --- | --- | --- | --- |
| Positive | 60 (80%) | 58 (77%) | 10 (13%) |
| Negative | 15 (20%) | 17 (23%) | 65 (87%) |

\*Significant difference between proportions using the Pearson Chi-square test

### SENV infection distribution, according to patient age, sex, and liver function indicators

Among patients with β-thalassemia, the highest frequencies of SENV-D (49.3%), SENV-H (37%), and combined SENV-D and SENV-H (13.7%) infections were observed in patients 15–26 years old (Table 2). Interestingly, SENV infections were not observed among any of the youngest donors.

**Table 2.**
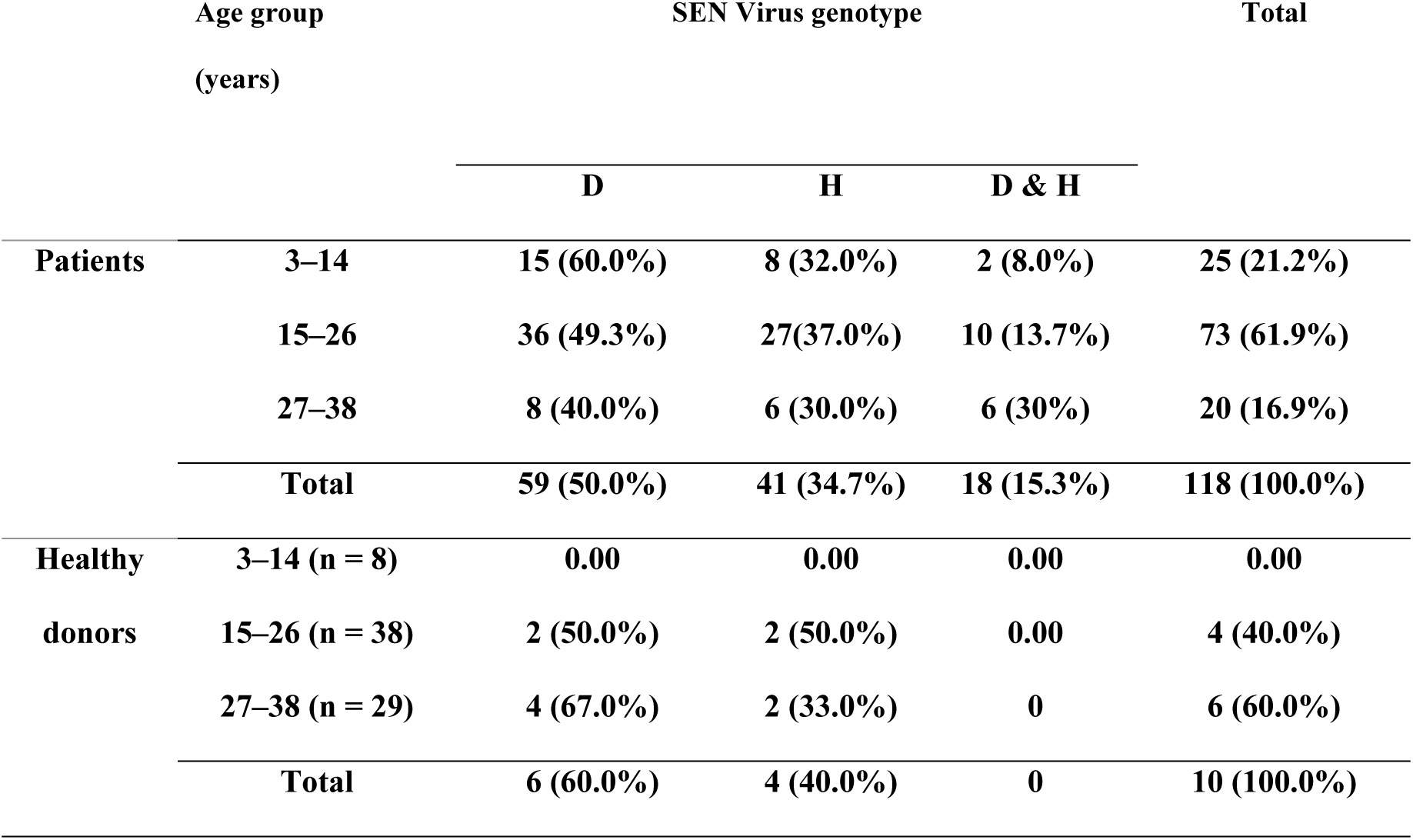
Age distribution of SEN virus infections among patients with β-thalassemia and healthy blood donors.

The sex-based distribution of SENV genotypes was also determined (Table 3). Among those with β-thalessemia, a high percentage of males (52%) were infected with SENV-D, whereas SENV-H was more predominant among females (40.3%). Co-infections with both SENV-D and SENV-H were most commonly observed in males (18.0%). However, a statistically significant sex-based distribution was not observed for either SENV genotype (Table 3).

**Table 3.**
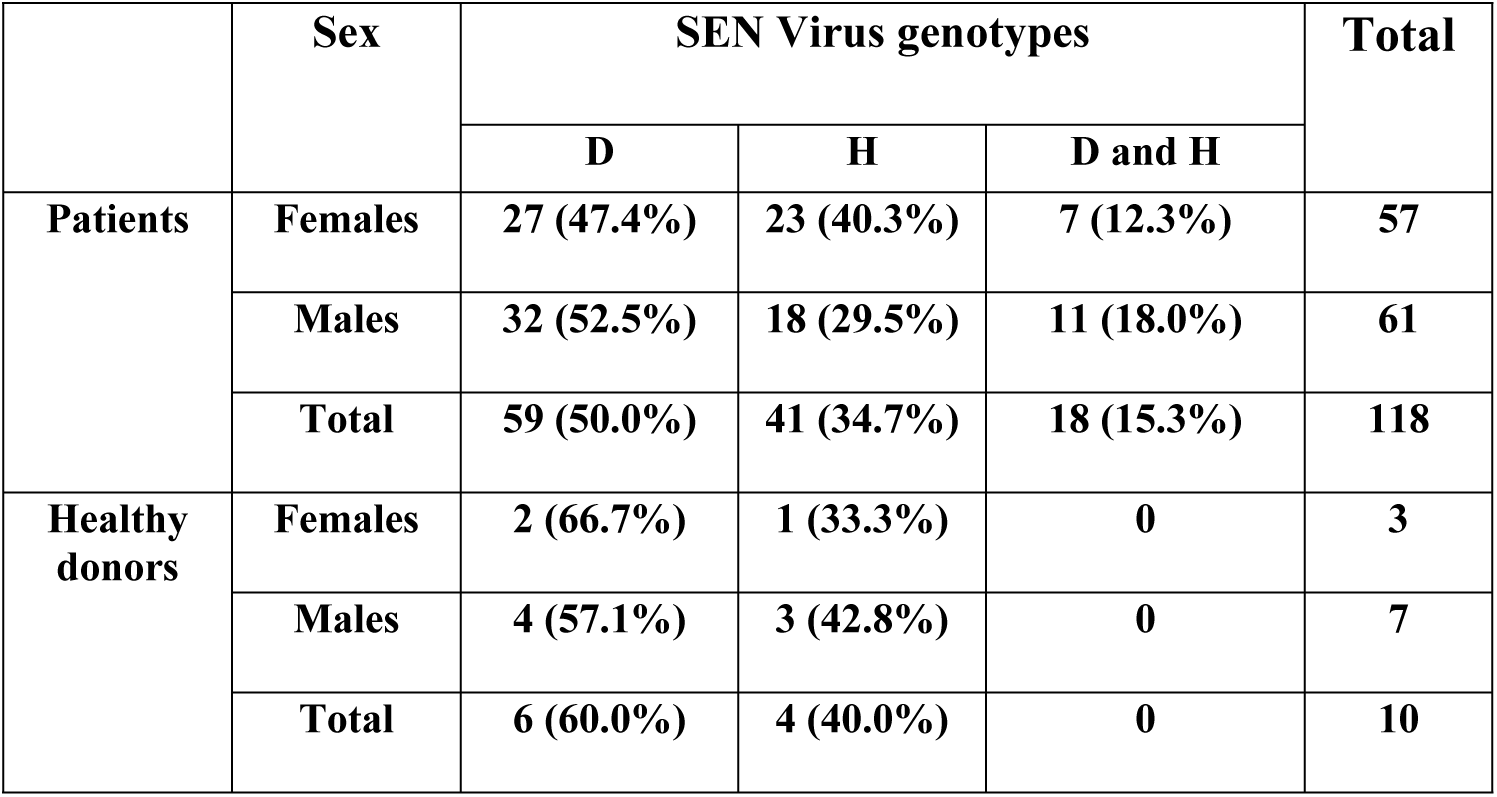
Sex-based distribution of SEN virus infections among patients with β-thalassemia and healthy blood donors.

|  | Sex | SEN Virus genotypes |  |  | Total |
| --- | --- | --- | --- | --- | --- |
|  |  | D | H | D and H |  |
| Patients | Females | 27 (47.4%) | 23 (40.3%) | 7 (12.3%) | 57 |
|  | Males | 32 (52.5%) | 18 (29.5%) | 11 (18.0%) | 61 |
|  | Total | 59 (50.0%) | 41 (34.7%) | 18 (15.3%) | 118 |
| Healthy donors | Females | 2 (66.7%) | 1 (33.3%) | 0 | 3 |
|  | Males | 4 (57.1%) | 3 (42.8%) | 0 | 7 |
|  | Total | 6 (60.0%) | 4 (40.0%) | 0 | 10 |

Significant differences were found in the activities of key liver function indicators in HCV-negative patients with SENV infections, relative to the healthy donors (Table 4). In the patients with SENV infections, most of the measured liver function indicators were also higher than the normal levels associated with healthy liver function. Moreover, the HCV-negative patients with thalassemia also showed liver function indicator (ALT, AST, ALP, TSB) levels that were significantly elevated compared with similar patients without SENV infections. Among the healthy donors, significant differences were not observed for any of the liver function indicators, regardless of SENV infection status.

**Table 4.** Liver function indicators (mean ± SD) in hepatitis C virus-negative patients with β-thalassemia with and without SEN virus infections and in healthy blood donors.

| Parameter | Patients with β-thalassemia | | P-value | Healthy donors | P-value |
| --- | --- | --- | --- | --- | --- |
|  | SENV-positive | SENV-negative |  |  |  |
| | Mean ± SD<br>(n = 58) | Mean ± SD<br>(n = 17) | | Mean ± SD<br>(n = 75) | |
| ALT (U/L) | 50.7 ± 13.6 | 30.7 ± 18.0 | <0.001 | 27.3 ± 7.02 | <0.001 |
| Normal: <40 |  |  |  |  |  |
| AST (U/L) | 50.8 ± 10.7 | 27.4 ± 10.5 | 0.001 | 24.2 ± 4.63 | <0.001 |
| Normal: <40 |  |  |  |  |  |
| ALP (U/L) | 135.9 ± 46.8 | 89.6 ± 17.1 | <0.001 | 71.1 ± 13.8 | <0.001 |
| Normal: <125 |  |  |  |  |  |
| GGT (U/L) | 50.1 ± 34.5 | 29.0 ± 6.5 | 0.229 | 27.6 ± 12.0 | <0.001 |
| Normal: <50 |  |  |  |  |  |
| TSB (mg/dL) | 2.7 ± 1.34 | 1.98 ± 1.07 | 0.001 | 0.99 ± 0.461 | <0.001 |
| Normal: <1.46 |  |  |  |  |  |

ALT, alanine transaminase; ASP, aspartate aminotransaminase; ALP, alkaline phosphatase; GGT, gamma-glutamyl transferase; TSB, total serum bilirubin

Comparing only HCV-negative patients with thalassemia to the healthy donors (Table 5), all liver function parameters, except GGT, were similarly elevated regardless of the genotype of the infecting SENV. This analysis revealed that GGT levels were significantly higher in patients with SENV-H infections compared with those observed in patients infected with SENV-D, combined SENV-D and H infections, or in healthy donors (all, P = 0.01). Small increases in other liver function parameter levels were also observed in the patients with SENV-H infections, relative to those with SENV-D infections (Table 5), but these differences were not statistically significant.

**Table 5.** Liver function marker levels (mean ± SD) in hepatitis C virus-negative patients with β-thalassemia infected with SEN virus and in healthy donors.

| Liver function marker | SEN Virus genotype |  |  |  | P-value |
| --- | --- | --- | --- | --- | --- |
|  | D<br>(n = 19) | H<br>(n = 31) | D and H<br>(n = 8) | Healthy donors<br>(n = 75) |  |
| ALT (IU/L) | 55.57 ± 18.5 | 60.44 ± 35.7 | 64.54 ± 16.4 | 38.7 ± 18.0 | <0.01 |
| AST (IU/L) | 52.21 ± 30.9 | 57.0 ± 17.6 | 54.3 ± 17.1 | 37.4 ± 15.5 | 0.03 |
| ALP (IU/L) | 134.4 ± 47.5 | 143.2 ± 54.0 | 135 ± 39.5 | 103.5 ± 57.1 | <0.01 |
| TSB (mg/dL) | 2.61 ± 1.32 | 2.87 ± 1.26 | 3.24 ± 1.38 | 1.7 ± 1.07 | 0.01 |
| GGT (IU/L) | 47.23 ± 28.6 | 85.2 ± 51.2 | 58.62 ± 18.3 | 37.6 ± 32.5 | 0.01 |

ALT, alanine transaminase; ASP, aspartate aminotransaminase; ALP, alkaline phosphatase; GGT, gamma-glutamyl transferase; TSB, total serum bilirubin

This study also showed significant differences in serum vitamin C, vitamin E, and GPX levels between healthy blood donors and SENV-positive, HCV-negative patients with thalassemia (Table 6). However, there was no significant difference in the levels of vitamins C and E between SENV-positive and SENV-negative patients. Interestingly, the level of GPX was significantly lower in SENV-positive patients than in SENV-negative patients.

**Table 6.** Vitamin C, vitamin E, and glutathione peroxidase levels in healthy blood donors and in SEN virus (SENV)-positive and -negative patients with β-thalassemia without hepatitis C virus infections.

| Parameter | Patients with β-thalassemia | | P-value | Healthy donors | P-value |
| --- | --- | --- | --- | --- | --- |
| | SENV-positive<br>Mean ± SD<br>(n = 58) | SENV-negative<br>Mean ± SD<br>(n = 17) | | (n = 75)<br>Mean ±SD | |
| Vitamin C<br>(μmol/L) | 58.9 ± 37.9 | 63.8 ± 40.1 | 0.871 | 73.0 ± 18.0 | 0.002 |
| Vitamin E<br>(μmol/L) | 32.4 ± 13.9 | 39.6 ± 15.1 | 0.628 | 44.6 ± 15.5 | <0.001 |
| Glutathione<br>peroxidase (U/L) | 238.2 ±<br>121.7 | 312.2 ±<br>127.3 | 0.049 | 353.5 ± 59.3 | <0.001 |

Although infection with SENV resulted in lower levels of these three antioxidants in the SENV-positive, HCV-negative thalassemia patients, compared with controls, only the GPX level appeared to be significantly lower in patients infected with SENV-H (P = 0.04). The distribution of antioxidants, including vitamins C and E and GPX, in patients with thalassemia and infected with the two SENV genotypes showed that the GPX levels were significantly lower in patients infected with SENV-D or SENV-H, than for healthy donors (Table 7).

**Table 7.** Glutathione peroxidase and vitamin C and E levels (mean ± SD) hepatitis C virus-negative patients infected with SEN virus and in healthy donors.

| Parameter | SEN V Genotypes |  |  |  | P-value |
| --- | --- | --- | --- | --- | --- |
|  | D<br>(n = 19) | H<br>(n = 31) | D and H<br>(n = 8) | Healthy<br>donors<br>(n = 75) |  |
| Vitamin C<br>(μmol/L) | 56.1 ± 36.9 | 58.6 ± 40.4 | 57.571 ± 38.6 | 62.85 ± 40.1 | 0.871 |
| Vitamin E | 31.7 ± 11.9 | 30.2 ± 14.4 | 37.7 ± 7.0 | 33.6 ± 15.1 | 0.628 |
| ($\mu\text{mol/L}$) | | | | | |
| **Glutathione peroxidase (U/L)** | $227.0 \pm 121.1$ | $217.1 \pm 116.8$ | $221.8 \pm 127.7$ | $282.6 \pm 127.3$ | 0.04 |

### Molecular and phylogenetic analyses

As part of this study, 14 samples of amplified SENV DNA were sent for sequencing and phylogenetic analyses, including 2 (14.3%) from healthy blood donors (nos. 15 and 17) infected with SENV-H, 9 (64.3%) from patients with thalassemia (nos. 1, 5–7, 10, 19, 20, 22, and 23) infected with SENV-H, and 3 (21.4%) from patients with thalassemia (nos. 2, 3, and 4) co-infected with SENV-D and SENV-H (Fig 3). An alignment study of SENV-H samples recovered from patients with thalassemia revealed a closely related genotype that is unique from samples isolated from patients in Iran, China, Japan, and France (GenBank accession numbers are documented in Table 8). Further, the sequencing of these genes revealed 85–97% compatibility with the global standard genes in GenBank.

**Table 8.** BLAST results of SEN virus genotype H DNA in GenBank, and DNA sequence compatibility with the global standard genes.

|  | Accession No. | Country | Source | Compatibility |
| --- | --- | --- | --- | --- |
| 1. | AY206683.1 | China | SEN virus strain SENV-H Orf2 gene | 93% |
| 2. | AB059353.1 | Japan | SEN virus SENV-H gene | 93% |
| 3. | KM593803.2 | France | SEN virus | 92% |
| 4. | KM593802.2 | France | SEN virus | 92% |
| 5. | GQ452053.1 | Iran | SEN virus Guilan SENV-H4 ORF1 gene | 97% |
| 6. | GQ452051.1 | Iran | SEN virus Guilan SENV-H2 ORF1 gene | 97% |
| 7. | AY183662.1 | China | SEN virus H gene | 85% |
| 8. | GQ179972.1 | Iran | SEN virus Guilan SENV-H1 ORF1 gene, | 94% |
| 9. | AB856068.1 | Iran | SEN virus SENV-H | 91% |
| 10. | AB856070.1 | Iran | SEN SENV-H | 90% |

**Fig 3.**
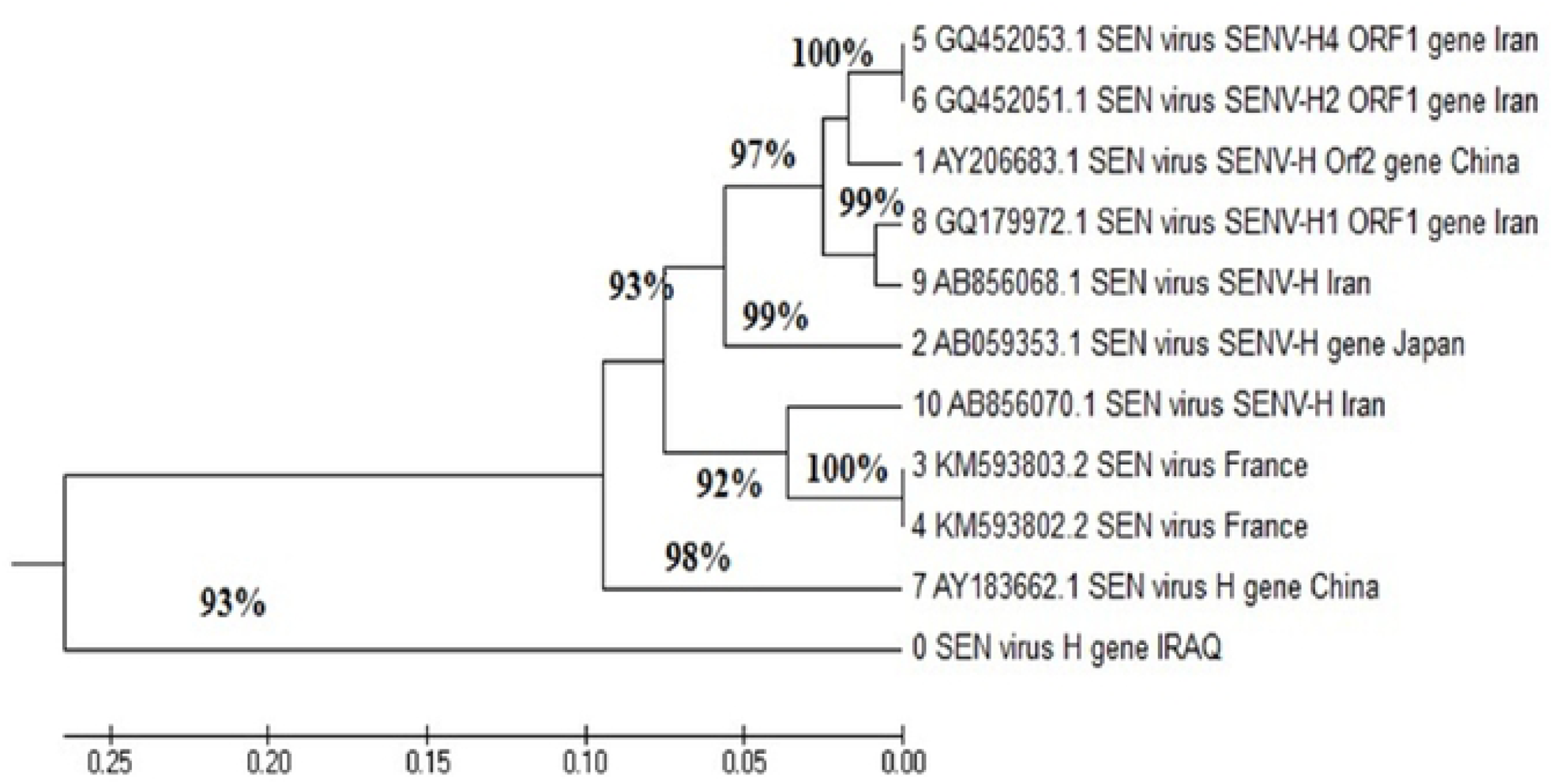
Phylogenetic tree analysis of the genetic distance between the Iraqi SEN virus H genotype and other sequences deposited in GenBank.

In the SENV-H gene sequencing study of isolates no. 1–7, 10, 15, 17, 19, 20, 22, and 23, the sequences corresponded to sequences already present in GenBank, specifically accession numbers: AY183662.1, KM593803.2, AY264849.1, AY206683.1, and AB059353.1. The sequencing results also revealed that after product alignment, the amplified SENV-H gene showed two types of substitutions (transition and transversion) compared with those in GenBank. Further, isolates 1 and 7 showed 100% identity with the sequence for AY183662.1. The remaining isolates yielded 92–99% identity; the low percentage of non-identity was due to transitions and transversions. The sequencing and BLAST analyses of the partial SENV-H genes and the types of gene polymorphisms are shown in Table 9.

**Table 9.**
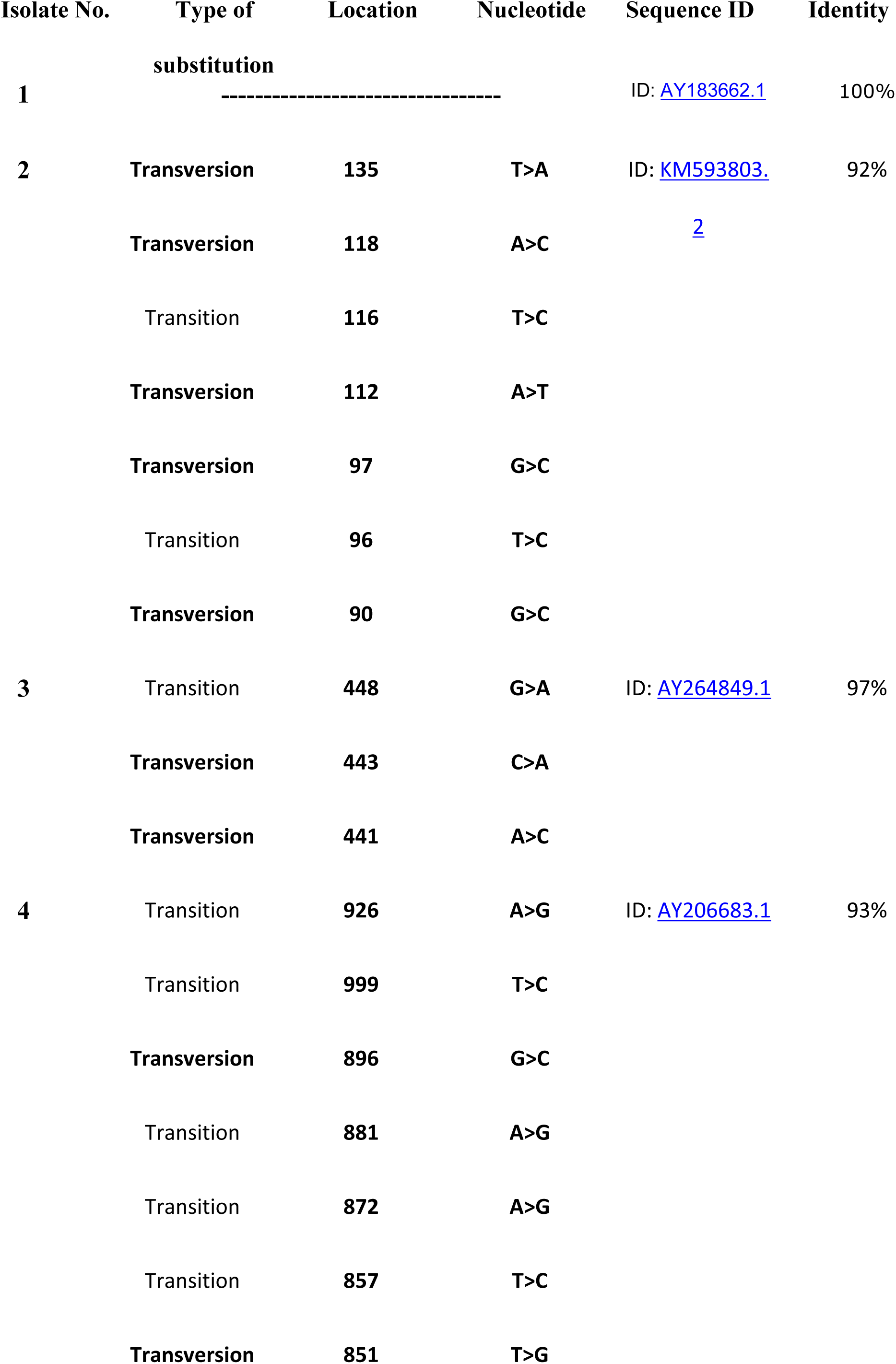
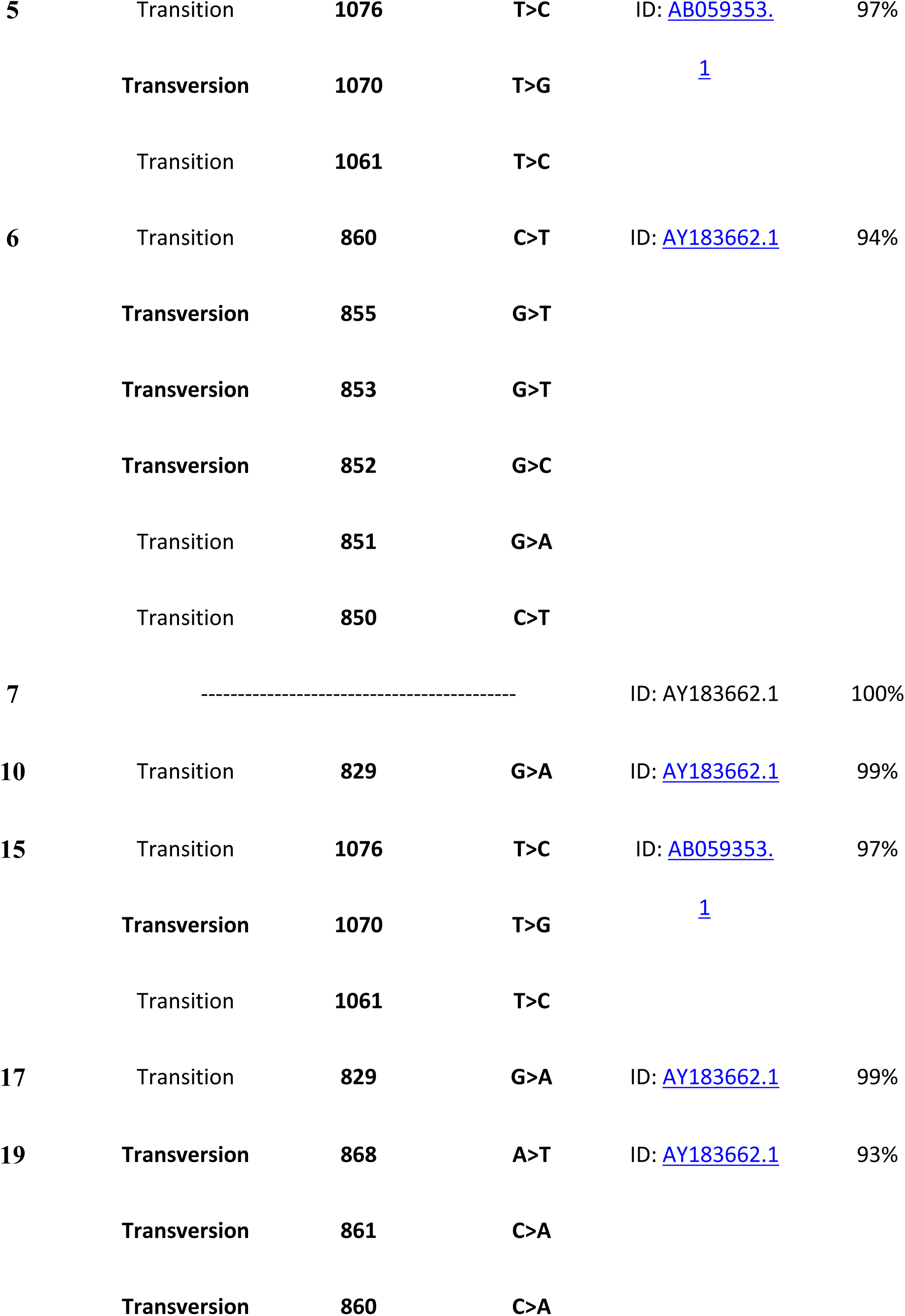
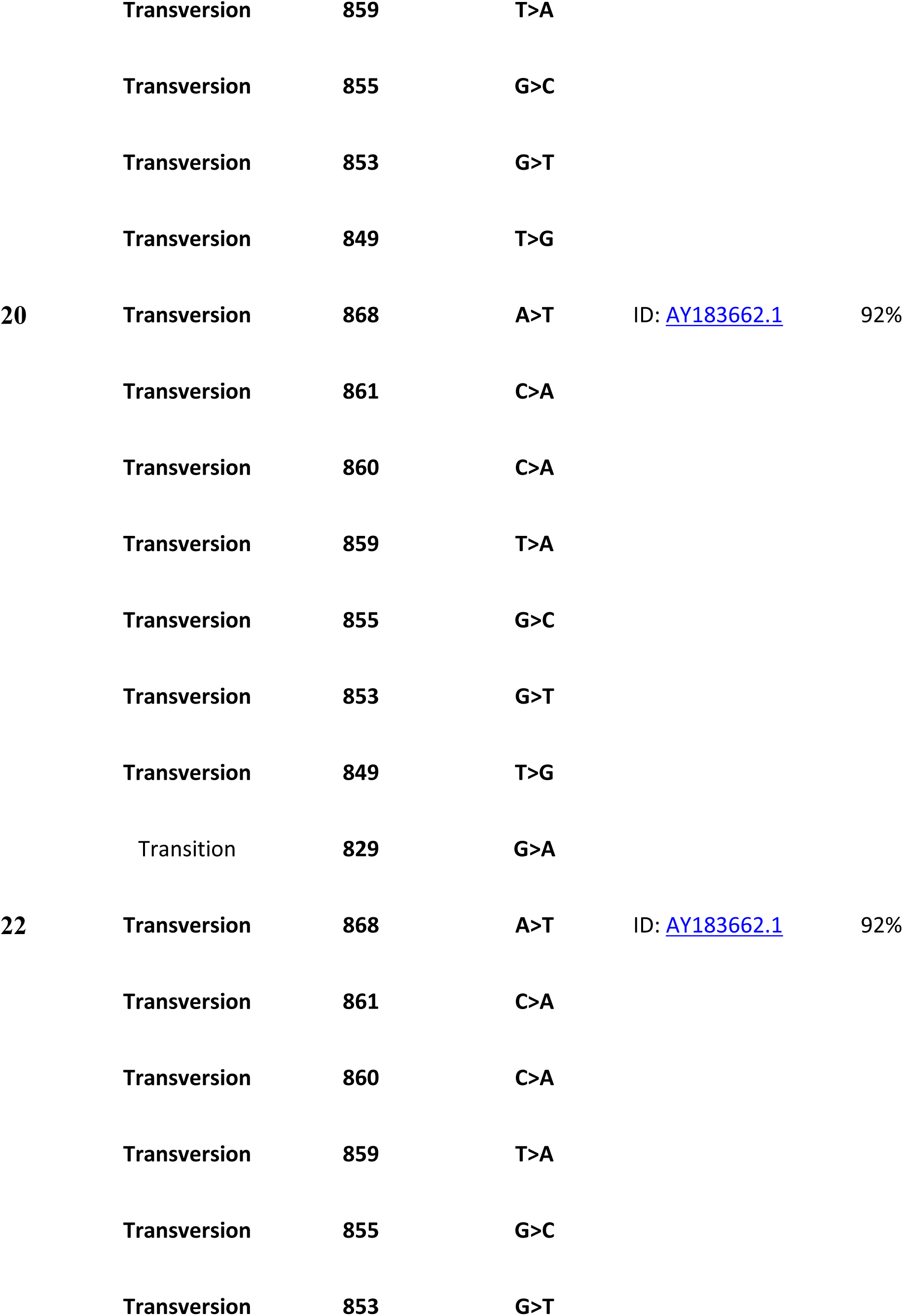
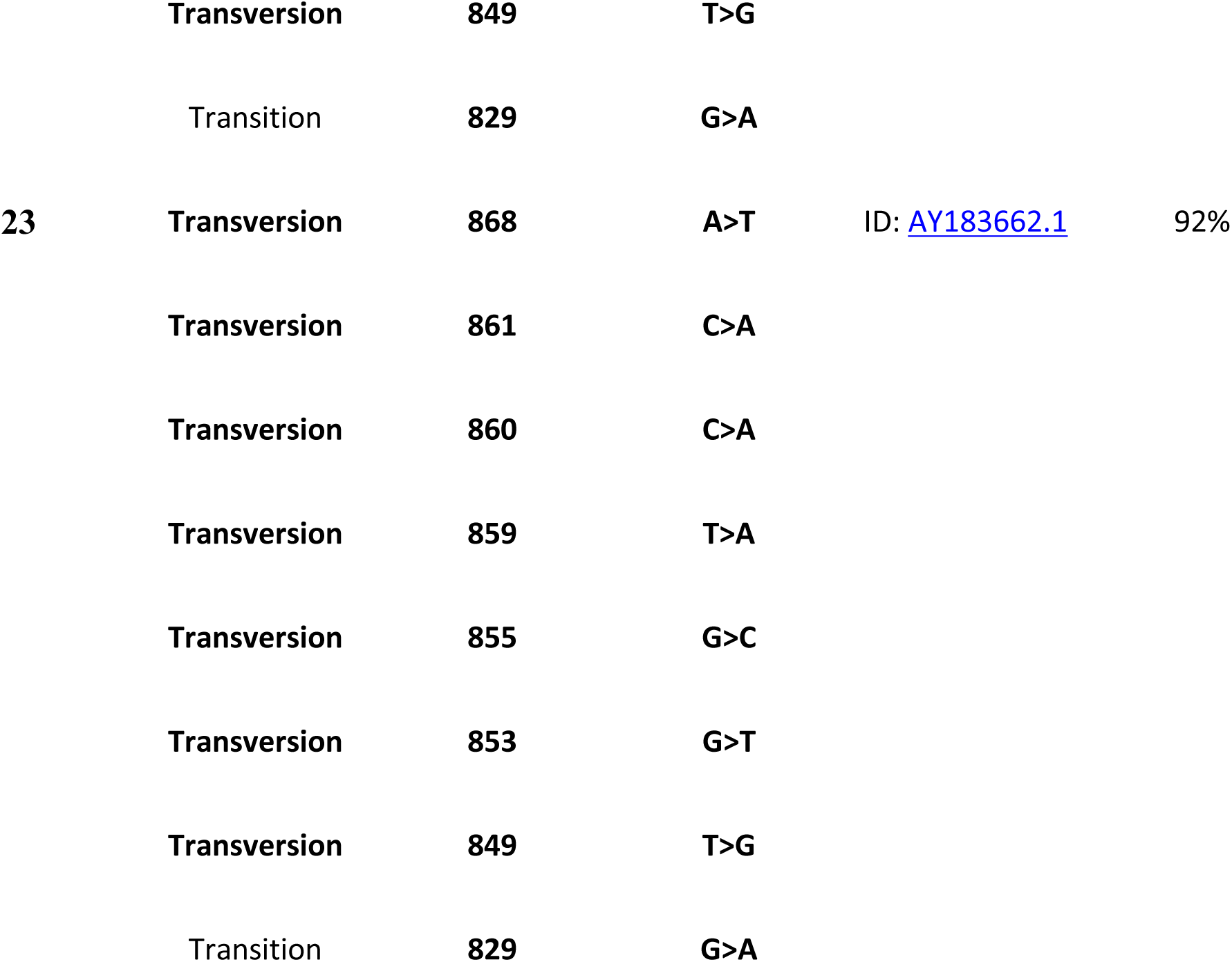
Characteristics of the clinical samples of SEN virus, genotype H.

## Discussion

The present study was a follow-up study from an earlier investigation of SENV infections in patients with β-thalassemia, in Iraq [8]. In addition to confirming the results from the earlier study, this study also sought to explore, in greater detail, the potential impact of SENV-H infections in patients with thalassemia. To this end, the study showed that the distribution of SENV-H is similar to that of SENV-D and that, generally, the physiological impacts of both genotypes of the virus are similar. However, the study also demonstrated that there are some differentiating hepatic effects, particularly related to GGT levels, in patients infected with SENV-H that are not observed in patients with SENV-D infections. SENV infections also appear to impact levels of some antioxidants in patients with adequate nutrition.

The present study confirmed that, in Iraq, SENV infections are more common in patients with thalassemia than in healthy donors. Further, patients with thalassemia demonstrated similar frequencies of SENV infection, regardless of their HCV infection status. The high rate of SENV infections among these patients reflects the similarly high rate of HCV infections typically observed in these patients [12]. The similarly high rates of infection for SENV and HCV in this patient population may be due to their having similar transmission risk factors, e.g., blood transfusions. Blood-borne infections, such as HCV, pose significant risks for patients with transfusion-dependent thalassemia [8]. Further, the significantly greater occurrence of SENV infections among patients with thalassemia than among healthy donors remains suggestive of blood transfusions being the primary source of SENV infections in patients with thalassemia [13]. Sani et al. [14] reported SENV viremia in 90.0% of patients with thalassemia (regardless of their HCV infection status) and in 76.7% of patients with HCV infections. These results are similar to the SENV infection rates reported in Taiwan [15]. Further, SENV infections in patients with thalassemia, regardless of their HCV status, have been suggested to further increase the patients need for blood transfusions [16]. This may be of particular concern since SENV transmission may occur primarily via parenteral routes, e.g., blood transfusion, intravenous drug use, or hemodialysis [17].

In this study, we also examined the age- and sex-based distributions of SENV infections. Among healthy donors, approximately 10.5% of 15–26-year-old donors were observed to have evidence of SENV infections as were 20.7% of 27–38-year-old donors. There was no evidence of a significant sex-based predominance. The prevalence and age distribution of SENV infections in this Iraqi population differs somewhat from an analysis of SENV infections in Taiwan, where 25–30% of adolescents were observed to have SENV infections [18]. Similar to the Taiwan study, however, there was a slight predominance of SENV-D infections compared with SENV-H infections. One major difference that was observed in the present study, compared with the Taiwan study, was that significantly more of the SENV infections in healthy Iraqi donors dwelt in urban areas. In the study of healthy adolescents in Taiwan [18], there were significantly higher rates of SEN viremia among those residing in rural or mountainous locales than for those living in urban environments. Regardless, SENV infections appear to occur in relatively young, healthy individuals, suggesting that the infection is unlikely to occur via a parenteral route in these individuals.

The SENV-H DNA samples recovered during the present study, from both patients with thalassemia and healthy donors, showed 97% compatibility with Iranian samples and 93% compatibility with samples from China. Moreover, the results showed 93% and 92% compatibility with samples from Japan and France, respectively. Samples in each branch of the SENV-H phylogenetic tree have corresponding DNA sequences, most probably indicating that the virus has been transmitted via blood transfusions. This is likely because of the DNA sequence compatibility between blood donors and patients with thalassemia and because the H sequences in isolates from individuals co-infected with both SENV-D and SENV-H are the same as the H sequences in isolates from individuals SENV-H infections. Karimi and Bouzari found a 90.8% frequency of SENV infections among healthy blood donors as well as high nucleotide homology in the sequenced amplicons of isolates from patients with thalassemia and healthy donors [4]. These results suggest that SENV-infected healthy blood donors act as partial sources of SENV transmission to patients with thalassemia and possibly to other individuals receiving transfusions. Karimi-Ratehkenari and Bouzari [16] found that a 90.8% frequency of SENV infection among healthy blood donors as well as high nucleotide homology of sequenced amplicons between isolates from patients with thalassemia. These results suggest that SENV-infected healthy blood donors act as partial sources of SENV transmission to patients with thalassemia and possibly to other individuals receiving transfusions. Therefore, our study suggested that the most probable mode of transmission to patients with thalassemia was through blood transfusions, based on the sequence homology between donors and recipients.

Our previous study showed that liver enzyme levels were significantly increased in thalassemia patients infected with HCV and/or SENV. Further, there appeared to be some additional increase associated in liver marker levels in patients co-infected with both viruses as opposed with just one or the other [8]. In the present study, we focused on the impacts of SENV infections in patients with thalassemia but who were not infected with HCV. The results of the current study showed that the liver function markers (ALT, ASP, AST, and TB) were elevated above normal values in HCV-negative patients with SENV infections and were significantly higher than in patients without either HCV or SENV infections or in healthy donors; the levels of the markers were similar between thalassemia patients not infected with either virus and healthy donors. Regardless, for most of the indicators of liver health, the elevations in marker levels were similar between patients infected with SENV-D and SENV-H. However, this was not the case for GGT. Among all HCV-negative patients with thalassemia, GGT levels were significantly elevated compared to the donors but were not significantly above normal values. However, when GGT levels were examined in patients infected with different genotypes of SENV, the levels were significantly higher in patients infected with SENV-H than in those infected with SENV-D. Moreover, the GGT levels were considerably above the normal range. Individuals infected with both SENV-D and SENV-H had GGT levels that were intermediate to the levels in patients infected with only SENV-D or SENV-H.

GGT is a known marker of oxidative stress because of its role in the catabolism of extracellular GPX (representative of intracellular antioxidants). A previous study showed that serum GGT levels are associated with clinical outcomes, even after adjusting for the presence of liver disease and liver function test results. Specifically, GGT levels are reportedly associated with the presence of HCC in patients with HCV infections [19]. Therefore, GGT levels might be predictive of HCC development in patients without cirrhosis, even after successful HCV eradication [20]. These observations suggest that a similar correlation might exist for patients with elevated GGT levels associated with SENV-H infections.

The association of GGT with oxidative stress prompted additional evaluations of the association of certain antioxidant levels with SENV infections, in the current study. Vitamins C and E both have antioxidant properties and were two of the antioxidants examined in this study. Vitamin C is a natural, water-soluble, free radical scavenger with the ability to donate two electrons from the double bond of its 6-carbon chain. During this process, oxidized vitamin C generates a stable intermediate product, dehydroascorbic acid, which can be taken up by erythrocytes and reduced to vitamin C via endogenous glutathione reductase [21]. In plasma, a relevant role of vitamin C is to restore α-tocopherol by oxidizing it. To be an effective antioxidant, the oxidized α-tocopherol must be reduced, but this process is slower than ascorbate recycling. Therefore, α-tocopherol is likely recycled in the cell membrane by a mechanism that involves enzymatic ascorbate recycling via α-tocopheroxyl [22]. To this end, some studies have suggested using dietary antioxidants, like vitamins C and E, to reduce liver enzyme levels in individuals with HCV infections [23]. In the present study, we found that the levels of vitamins C and E were significantly lower in thalassemia patients than in the healthy donors. However, the levels were similar between patients with and without SENV infections. Additionally, there was no difference in vitamin C/E levels between patients infected with either genotype of SENV. there were also no differences in the levels of vitamins C and E associated with SENV-D or SENV-H infections. When we examined the levels of GPX in the same individuals, the levels were significantly lower in the thalassemia patients than in the healthy donors. Further examination also revealed that the levels of GPX were lower in patients infected with SENV-H than in those infected with SENV-D.

This study is subject to some limitations. First, the retrospective design of the study makes it susceptible to the same type of bias that is common to all retrospective studies, particularly since consecutive patients were recruited for the study based on a clinical diagnosis that was suggestive of the presence of HCV infection. Second, the study population was relatively small. Additional studies will be needed to confirm the accuracy of the conclusions derived from the present study. Finally, since this study was limited to a specific area of Iraq, the ability to generalize the results to other geographic areas and ethnic groups of people is limited.

In conclusion, this study confirmed the results of our previous study that suggested that, in Iraq, the prevalence of SENV infections is higher in patients with β-thalassemia than in healthy blood donors, but that the presence of HCV infections does not affect the prevalence of SENV infections. Further, this study was able to demonstrate the absence of a sex-based predominance of infection by the SENV D and H genotypes. The study also showed that, in the present population, the SENV-H DNA indicated a high degree of similarity with other global isolates deposited in GenBank, especially with those from Iran. SENV-H was also suggested to play a more pronounced role in raising serum levels of GGT. Further, SENV infections were shown to depress levels of GPX to a greater extent than in patients without SENV infections.

## Competing Interests

No competing interests exist.

## Acknowledgement

The authors acknowledge Harkynn Consulting (www.harkynn.com) for English language editing assistance.

## References

1. Kadhim KA, Baldawi KH, Lami FH. Prevalence, incidence, trend, and complications of thalassemia in Iraq. Hemoglobin. 2017;41: 164–168. doi: 10.1080/03630269.2017.1354877.

2. Chethan C, Valliyamma V. Correlation study between serum AFP an ALT (SGPT) level in acute viral hepatitis. Acta Biomedica Scientia. 2017;4 11–13. doi: 10.21276/abs.2017.4.1.2.

3. Umemura T, Yeo AE, Sottini A, Moratto D, Tanaka Y, Wang RY, et al. SEN virus infection and its relationship to transfusion-associated hepatitis. Hepatology. 2001;33: 1303–1311. doi: 10.1053/jhep.2001.24268.

4. Tanaka Y, Primi D, Wang R, Umemura T, Yeo A, Mizokami M, et al. Genomic and molecular evolutionary analysis of a newly identified infectious agent (SEN virus) and its relationship to the TT virus family. J Infect Dis. 2001;183: 359–367. doi: 10.1086/318091.

5. Yoshida EM, Wong SG. SEN virus. A clinical review. Minerva Gastroenterol Dietol. 2002;48: 73–79.

6. Akiba J, Umemura T, Alter HJ, Kojiro M, Tabor E. SEN virus: epidemiology and characteristics of a transfusion-transmitted virus. Transfusion. 2005;45: 1084–1088. doi: 10.1111/j.1537-2995.2004.00209.x

7. Ghasemi-Dehkordi P, Doosti A. The prevalence of SEN virus infection in blood donors and chronic hepatitis B and C patients in Chaharmahal Va Bakhtiari province. Journal of Cell and Animal Biology. 2011;5: 182–186.

8. Al-Ani SK, Al-Ouqaili MTS, Awad MM. Molecular and genotypic study of SENV-D virus co-infection in β-thalassemic patients infected with the hepatitis C virus in Iraq. International Journal of Green Pharmacy. 2018;12(4 Suppl): 1–11. doi: 10.22377/ijgp.v12i04.2276.

9. Al-Ouqaili MTS, Al-Hayani NN, Saadoon IH. Detection of rhinovirus and some DNA/RNA viruses by reverse transcriptase real-time PCR and their immunological parameters in patients with acute respiratory tract infection in Iraq: Molecular and immunological study. International Journal of Pharmaceutical Sciences and Research. 2019;10: 1000–1011. doi: 10.13040/IJPSR.0975-8232.10(5).1000-11.

10. Al-Ouqaili MTS, Al-Kubaisi SHM. Molecular genetics study on high and intermediate risk genotypes of human papillomavirus among patients with benign and malignant cervical lesions. International Journal of Pharmaceutical Sciences and Research. 2019;10: 1000–1009. doi: 10.13040/IJPSR.0975-8232.10(4).1000-09.

11. Tamura K, Stecher G, Peterson D, Filipski A, Kumar S. MEGA6: Molecular evolutionary genetics analysis version 6.0. Mol Biol Evol. 2013;30: 2725–2759 doi: 10.1093/molbev/mst197.

12. Bhattacharyya KK, Biswas A, Gupta D, Sadhukhan PC. Experience of hepatitis C virus seroprevalence and its genomic diversity among transfusion-dependent thalassemia patients in a transfusion center. Asian Journal of Transfusion Science. 2018;12: 112–116.

13. Abbasi S, Makvandi M, Karimi G, Neisi N. The prevalence of SEN virus and occult hepatitis B (OBI) virus infection among blood donors in Ahvaz City. Jundishapur J Microbiol. 2016: 9: e37329. doi: 10.5812/jjm.37329

14. Sani HO, Sharifi Z, Hosseini SM, Shooshtari MM. SEN virus detection in thalassemic patients infected with hepatitis C virus. Arch Virol. 2012;157: 2441–2445. doi: 10.1007/s00705-012-1443-3.

15. Kao JH, Chen W, Chen PJ, Lai MY, Chen DS. Prevalence and implication of a newly identified infectious agent (SEN virus) in Taiwan. J Infect Dis. 2002;185: 389–392. doi: 10.1086/338472.

16. Karimi-Rastehkenari A, Bouzari M. High frequency of SEN virus infection in thalassemic patients and healthy blood donors in Iran. Virol J. 2010;7:1 doi: 10.1186/1743-422X-7-1.

17. Hosseini SA, Bouzari M. Detection of SENV virus in healthy, hepatitis B- and hepatitis C-infected individuals in Yazd Province, Iran. Iran Biomed J. 2016;20:168–174.

18. Wang LY, Ho TY, Chen MC, Yi CS, Hu CT, Lin HH. Prevalence and determinants of SENV viremia among adolescents in an endemic area of chronic liver diseases. J Gastroenterol Hepatol. 2007;22: 171–176. doi: 10.1111/j.1440-1746.2006.04537.x.

19. Huang CF, Yeh ML, Tsai PC, Hsieh MH, Yang HL, Hsieh MY, et al. Baseline gamma-glutamyl transferase levels strongly correlate with hepatocellular carcinoma development in non-cirrhotic patients with successful hepatitis C virus eradication. J Hepatol. 2014;61: 67–74. doi: 10.1016/j.jhep.2014.02.022.

20. Huang R, Yang CC, Liu Y, Xia J, Su R, Xiong YL, et al. Association of serum gamma-glutamyl transferase with treatment outcome in chronic hepatitis B patients. World J Gastroenterol. 2015;21: 9957–9965. doi: 10.3748/wjg.v21.i34.9957.

21. Nimse SB, Pal D. Free radicals, natural antioxidants, and their reaction mechanisms. RSC Advances. 2015;5(35):27986–28006. doi: 10.1039/C4RA13315C.

22. Gonçalves D, Lima C, Ferreira P, Costa P, Costa A, Figueiredo W, et al. Orange juice as a dietary source of antioxidants for patients with hepatitis C under antiviral therapy. Food Nutr Res. 2017;61:1296675. doi: 10.1080/16546628.2017.1296675.

23. Ji HF, Sun Y, Shen L. Effect of vitamin E supplementation on aminotransferase levels in patients with NAFLD, NASH, and CHC: results from a meta-analysis. Nutrition. 2014;30: 986–991 doi: 10.1016/j.nut.2014.01.016.

